# Comprehensive analysis of RNA-seq and whole genome sequencing data reveals no evidence for SARS-CoV-2 integrating into host genome

**DOI:** 10.1101/2021.06.06.447293

**Authors:** Yu-Sheng Chen, Shuaiyao Lu, Bing Zhang, Tingfu Du, Wen-Jie Li, Meng Lei, Yanan Zhou, Yong Zhang, Penghui Liu, Yong-Qiao Sun, Yong-Liang Zhao, Ying Yang, Xiaozhong Peng, Yun-Gui Yang

## Abstract

SARS-CoV-2, as the causation of severe epidemic of COVID-19, is one kind of positive single-stranded RNA virus with high transmissibility. However, whether or not SARS-CoV-2 can integrate into host genome needs thorough investigation. Here, we performed both RNA sequencing (RNA-seq) and whole genome sequencing on SARS-CoV-2 infected human and monkey cells, and investigated the presence of host-virus chimeric events. Through RNA-seq, we did detect the chimeric host-virus reads in the infected cells. But further analysis using mixed libraries of infected cells and uninfected zebrafish embryos demonstrated that these reads are falsely generated during library construction. In support, whole genome sequencing also didn’t identify the existence of chimeric reads in their corresponding regions. Therefore, the evidence for SARS-CoV-2’s integration into host genome is lacking.

**One-Sentence Summary:** SARS-CoV-2 does not integrate into host genome through whole genome sequencing.

---

Severe Acute Respiratory Syndrome Coronavirus 2 (SARS-CoV-2) is an RNA virus of the Coronaviridae family causing the outbreak and worldwide pandemic of Coronavirus Disease 2019 (COVID-19)(*1–3*). It has become a public health emergency and an unprecedented global threat affecting about 162 million individuals leading to over 3.3 million deaths up to the middle of May, 2021. In attributed to the extensive studies on SARS-CoV-2, several kinds of vaccines are currently available which brings hope to human society for alleviating and eventually preventing COVID-19 epidemic (*4–6*). However, a better understanding of viral pathogenesis, particularly the viral-host interaction, are needed to develop effective interventions.

Viruses are mainly separated into four groups based on the types of genome they belong to, including dsDNA, ssDNA, dsRNA and ssRNA (*7*). All viruses need host cells for their reproduction through introducing their genetic materials into the infected cells and hijacking certain cellular machinery, such as ribosome, polymerase and so on, to produce viral particles (*8–13*). Noticeable, some RNA viruses are capable of inserting their reverse-transcribed DNA into host genome and then produce viral RNA through host-dependent transcription pathway. To fulfill this, two mechanisms have been reported: (1) some retroviruses, such as HIV (human immunodeficiency virus), ASV (avian sarcoma virus) and PFV (prototype foamy virus), can encode their own reverse transcriptase to synthesize a complementary ssDNA for the host genome integration (*14, 15*); (2) for some non-retroviruses, such as LCMV (lymphocytic choriomeningitis virus) and BDV (borna disease virus), they have been reported to utilize elements of endogenic transposons, like endogenous retroviral transposon IAP (intracisternal A-particle) or LINE-1 (long interspersed element-1), for recombination with host genome (*16*). It should be noted that the endogenous retrotransposons are inactivated in mammalian cells except for early developmental stage or pathological conditions, such as tumors (*17*).

As a positive single-strand RNA virus, SARS-CoV-2 generates sub-genomic RNAs through a mechanism termed discontinuous extension of minus strands, and further synthesizes its proteins in cytoplasm of infected cells (*18*). However, one recent study revealed that SARS-CoV-2 infection induced retrotransposon activation, leading to the formation of chimeric virus-retrotransposon RNA (*19*). Zhang et al. also observed the host genome integration of SARS-CoV-2 under overexpressing LINE-1 in cultured human cells (*20*). These reports indicate a likely chance of SARS-CoV-2 integration into host genome (*19, 20*), which have fueled concerns about its potential long-term health threat in the infected individuals. Thus, more evidences are explicitly required to address this issue. Here, we performed both RNA sequencing and whole genome sequencing in samples from SARS-CoV-2 infected human and monkey cells, and investigated the presence of host-virus chimeric events. We found that the chimeric reads from RNA-seq were artificially contributed by library construction. Therefore, no evidence from our study supports the likely integration of SARS-CoV-2 into host genomes.

To investigate whether or not the SARS-CoV-2 is capable of integrating into host genome, we performed RNA-seq for SARS-CoV-2 infected 293T, Huh-7 and Calu-3 cells and validated the chimeric reads from both SARS-CoV-2 and human genomes. Firstly, we assessed the reproducibility of chimeric junctions in one sample or amongst three samples. CPM (Counts Per Million) was estimated by the counts of all chimeric reads along gene body and normalized by sequencing depth (fig. S1A). After filtering genes with low chimeric levels, there were 126 chimeric genes identified in Huh-7 and Calu-3 cells, while none was detected in 293T cells. Moreover, 18 of them were overlapped between these two cell lines, and covered various RNA types, including mRNA, lncRNA and miscellaneous RNA (misc RNA) (fig. S1B). We further checked the proportion of reads derived from SARS-CoV-2 in each cell line and found that only 0.4% reads were aligned to viral genome in 293T cells, much less than that of the other two cell lines (fig. S1C), which is supported by read counts for all genes containing chimeric reads (fig. S2A). As such, we used the viral depth, instead of sequencing depth, for CPM normalization.

Using the normalized CPM, there were 347, 3107 and 4171 chimeric genes identified from 293T, Huh-7 and Calu-3 cells, respectively (Fig. 1A and fig. S2B). Most of them were the fusion of viral RNA with mRNA from host cells (fig. S2B). We further obtained 132 conserved chimeric genes from all of three cell lines (Fig. 1B) and analyzed the categories of RNA types. The results showed that the chimeric genes were preferentially mRNA (Fig. 1C), likely due to the abundant expression of mRNA relative to other RNA types. We then displayed the chimeric events for 5 randomly picked conserved chimeric genes (*RPL3, GPI, ATP5F1A, EEF2* and *RPS19*) among 293T, Huh-7 and Calu-3 cell lines, and found that the chimeric events were extremely diverse in these cells (Fig. 1D and fig. S2C). Moreover, we analyzed the number of each chimeric event for chimeric gene, and found that most of them could be identified only once in each cell line (fig. S2D). Collectively, these results suggest that RNA chimeric event seems to be a random case for an individual infected cell.

**Fig. 1.**
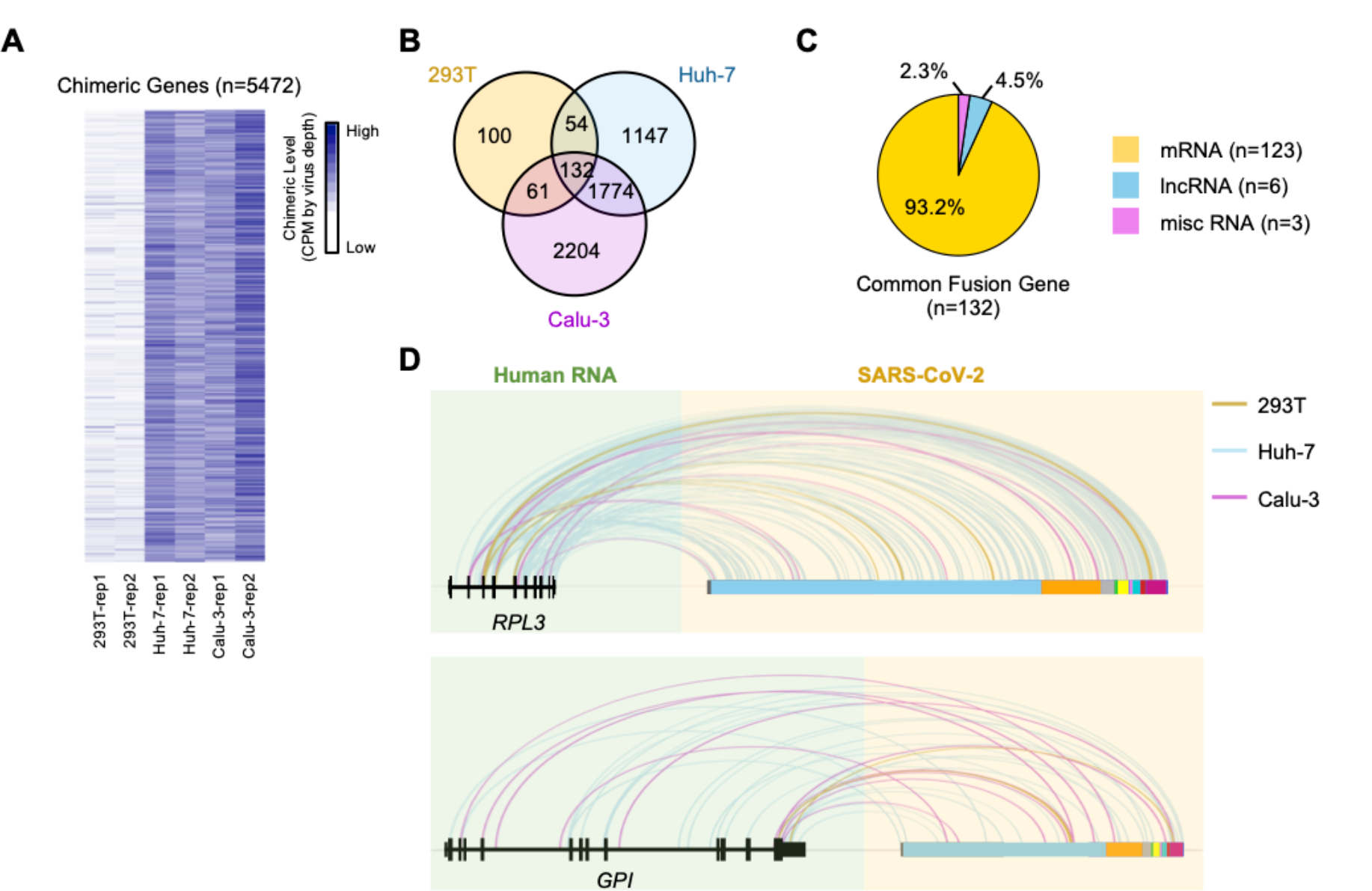
Chimeric genes of human and SARS-CoV-2 were detected by RNA-seq. (**A**) Heatmap displaying the chimeric level for filtered chimeric genes in each sample. The chimeric level was defined as CPM of chimeric reads and normalized by depth of reads from SARS-CoV-2. (**B**) Venn diagram showing the number of shared and specific chimeric genes among 3 human cell lines. (**C**) Pie chart displaying proportion of RNA types for 132 common chimeric genes. (**D**) Integrative Genomics Viewer (IGV) tracks displaying the junction loci in both human RNA and SARS-CoV-2 RNA for common chimeric genes *RPL3* (upper) and *GPI* (lower). The lines with different colors indicate the sources of chimeric reads, where yellow, blue and pink represents 293T, Huh-7 and Calu-3 cells, respectively. Blocks with different colors represent 5’UTR, ORF1ab, S, ORF3a, E, M, ORF6, ORF7ab, ORF8, N and 3’UTR along the SARS-CoV-2 genome.

To further investigate whether chimeric events were species-specific, we performed RNA-seq on SARS-CoV-2 infected green monkey Cos-7 and rhesus monkey MA-104 cells. Intriguingly, compared to Cos-7, there was a large amount of viral RNA detected in MA-104 sample (fig. S3A). Moreover, most chimeric genes estimated by CPM contained only one read and belonged to mRNA (fig. S3,B and C). Similar to human cell line, the chimeric events could only be found once in each monkey cell line (fig. S3D). Thus, low incidence of chimeric event was observed in the infected cells from human and other species.

We next tried to determine whether the diversity of chimeric events among different cell types results from random integration or integration preference for specific cell types, we displayed the overlapped chimeric genes and events for each two cell lines (Fig. 2A). Though over half chimeric genes overlapped in both cell lines, the shared chimeric events were extremely low. We then used different resolutions to define the chimeric events to determine if the virus-host integration might be enriched in hot-spot regions instead of specific nucleotides, and found these events were still not conserved (Fig. 2A). Given that the low reproducibility might be contributed by heterogeneity from different cell lines, we further analyzed the number of overlapped chimeric genes and events between two sequencing replicates from one cell line (fig. S4A). Unexpectedly, the chimeric genes in 293T cells were diverse with only few chimeric events identified in both replicates (fig. S4, A and B), which might be owing to the much less SARS-CoV-2 reads in this cell line.

**Fig. 2.**
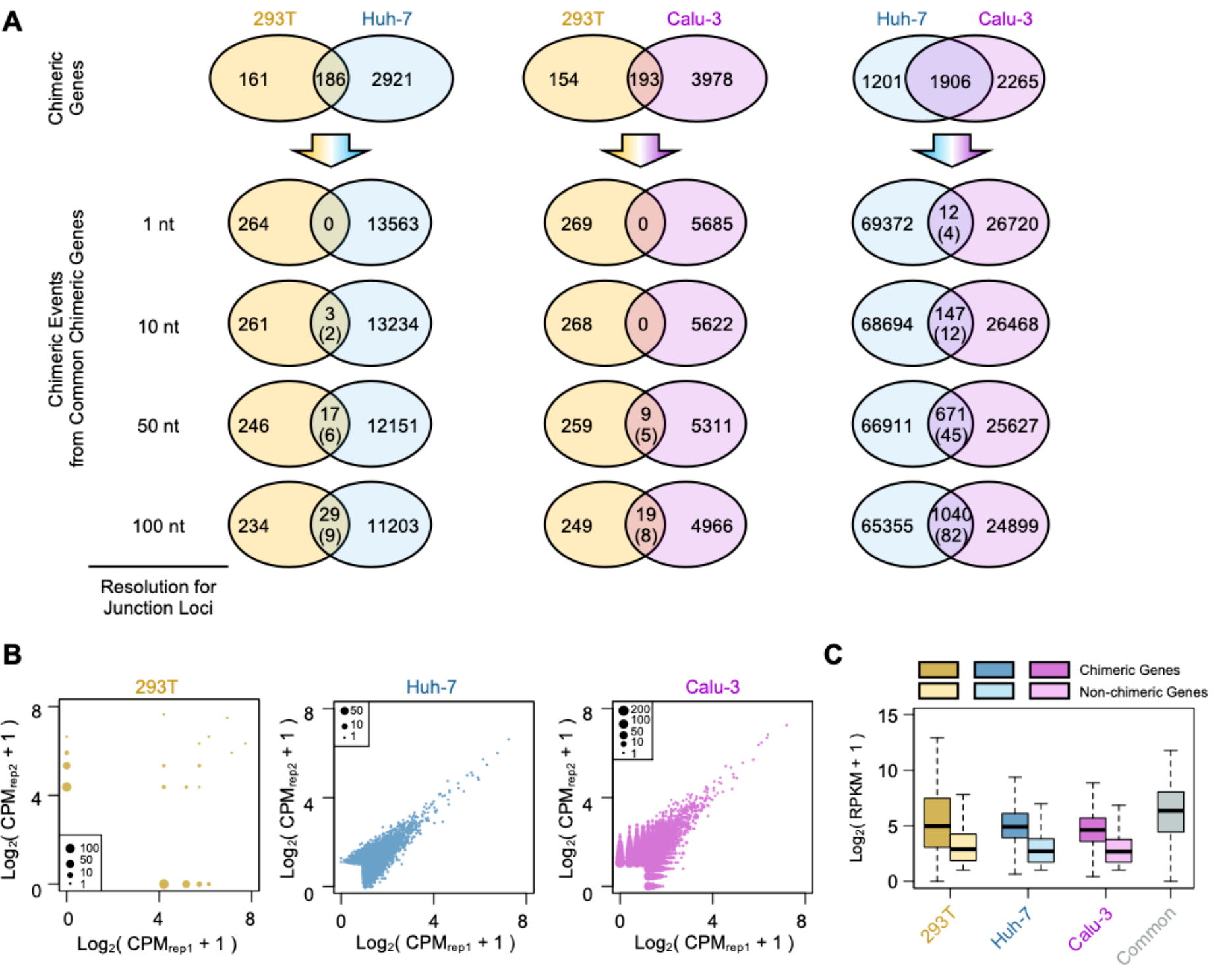
Chimeric genes were frequently detected among highly expressed genes. (**A**) Venn diagrams showing the number of shared chimeric genes for each two cell lines (top) and conserved chimeric events on shared chimeric genes at different resolutions (1 nt, 10 nt, 50 nt and 100 nt). For venn diagrams of chimeric events, the value in parenthesis represents the number of chimeric genes containing the shared chimeric events. (**B**) Scatter plot displaying the chimeric levels for chimeric genes in two replicates. The sizes of scatters represent number of chimeric genes with the same distribution of chimeric levels in two replicates. (**C**) Boxplot displaying the distributions of expression level for chimeric genes (dark color) and non-chimeric genes (light color) in each cell line. The grey block represents expression levels of 132 common chimeric genes.

Then we compared the chimeric level for each chimeric gene in two replicates (Fig. 2B). The relatively high similarity between two replicates was found in both Huh-7 and Calu-3 cells, suggesting that SARS-CoV-2 might select genes to fuse by some unknow elements in individual cells while the loci seemed random. To further search for the key elements, we compared the expression levels between chimeric and non-chimeric genes, and observed that chimeric genes had a remarkably higher expression level than those of non-chimeric genes (Fig. 2C and fig. S5A). After evaluating the proportions of chimeric genes with low to high expression level, chimeric events tend to occur in highly expressed host genes (fig. S5B). In addition, the chimeric loci along virus were also enriched in N region, the most abundant sub-genomic RNA of SARS-CoV-2 (fig. S6, A and B), indicating that chimeric events were positively correlated with expression levels of both host and virus RNAs.

We next tested the conservativeness of the preference of chimeric events for highly expressed genes in monkey cell lines. The reproducibility for two replicates were first validated for chimeric genes (fig. S7A) and the chimeric events for the shared chimeric genes were then analyzed (fig. S7, B and C). Similar to the findings in human cells, reproducibility of chimeric events in overlapped genes were extremely low between two replicates, suggesting that the integrating events were most likely randomly occurred. In addition, chimeric genes showed preferred enrichment in high expression genes in both monkey cell lines (fig. S7, D-F). Moreover, the viral chimeric loci were highly correlated to the expression of viral sub-genomic RNA in MA-104 (fig. S7, G and H), which was also observed in human cell lines.

Recently, Zhang et al. reported that SARS-CoV-2 RNA might integrate into host genome *via* reverse-transcription, and overexpressed LINE-1 might stimulate the reverse-transcribed SARS-CoV-2 integration into host genome (*20*), which prompted us to determine whether or not the correlation of chimeric events with high expression of host genes was attributed to the viral integration into the locations of DNA elements, such as promoters and enhancers. We first analyzed the number of chimeric genes in each chromosome, and didn’t identify any evidence of preferred viral integration into some chromosomes (fig. S8, A and B).

We then estimated the accumulated chimeric and expression levels of 100 genes in each of 617 bins obtained by sequentially cutting the genome along chromosomes (each containing 100 genes). Intriguingly, though there was no preference for whole chromosomes, the bins represented positive correlation between expression and chimeric levels (Fig. 3A), which might be explained by possible viral integration into transcription regular elements along host DNA. Additionally, we found that the bin in chromosome 14 (chr14: 39,385,404-49,852,821) had both the highest chimeric level and gene expression, and was well conserved among 3 cell lines (Fig. 3, A and B). However, this much enhanced chimeric level was likely to be contributed by one single gene, the non-coding *RN7SL1*, with the highest expression. Next, we tried to use the frequency of chimeric genes to explain whether the chimeric preference to highly expressed genes resulted from integration (fig. S8, C and D). However, no evidence could support this hypothesis.

**Fig. 3.**
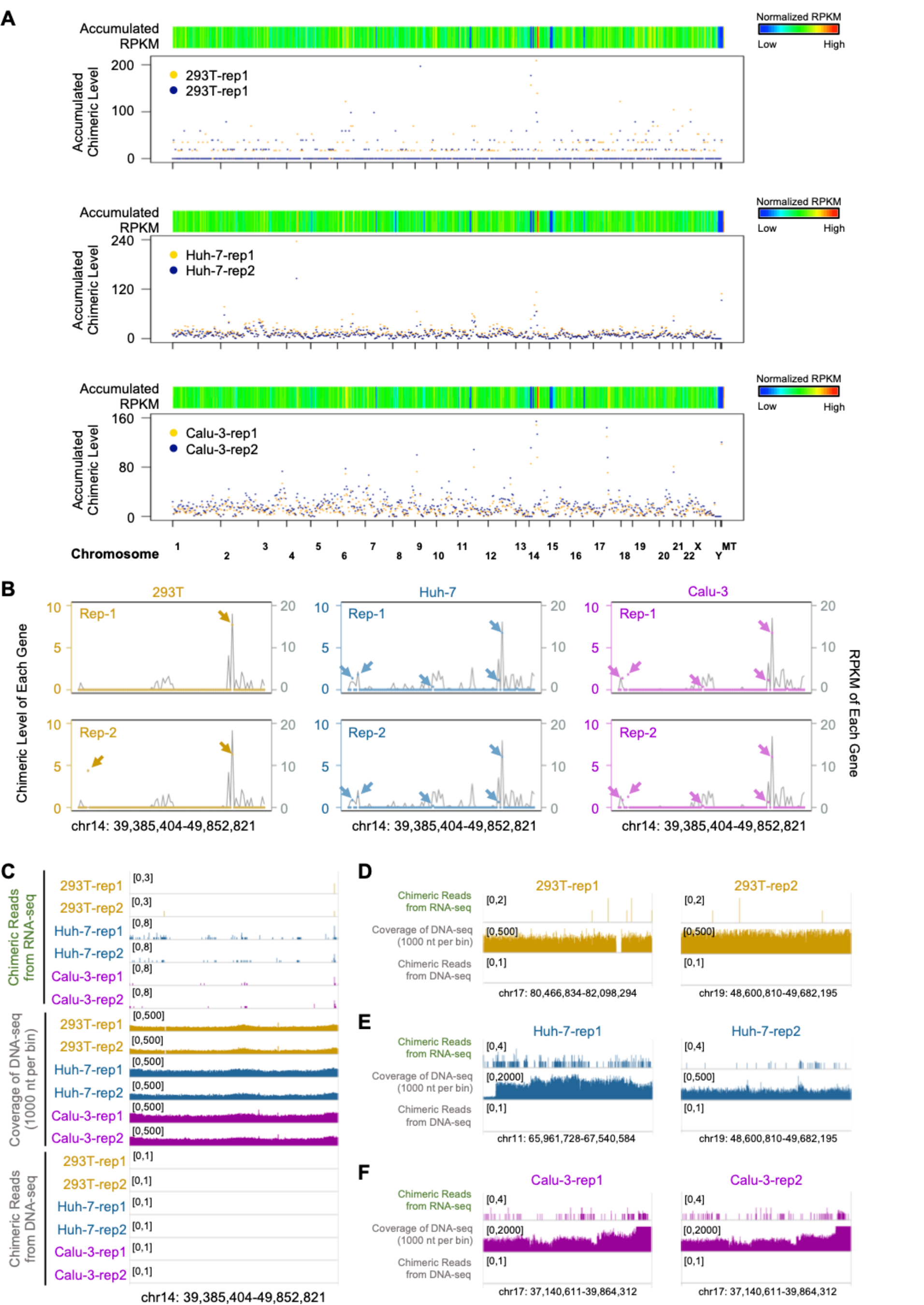
Whole genome sequencing reveals no integration of SARS-CoV-2 into host genome. (**A**) Heatmap showed the accumulated expression score of each bin along human genome, and scatter plot below displayed the accumulated chimeric levels of corresponding bins for each cell line. (**B**) Chimeric levels and corresponding expression levels of genes along the bin with the highest accumulated chimeric levels (chr14: 39,385,404-49,852,821) were displayed by scatters and lines, respectively. The arrows point out the chimeric levels and gene loci of chimeric genes. (**C**) IGV tracks displaying the expression level (top), coverage of whole genome sequencing (middle) and genomic insertion signal (bottom) along chr14: 39,385,404-49,852,821. No viral integrating signal could be detected among all samples. (**D to F**) IGV tracks displaying the expression level (top), coverage of whole genome sequencing (middle) and genomic insertion signal (bottom) along bins in 293T (**D**), Huh-7 (**E**) and Calu-3 (**F**).

To determine whether or not the viral integration directly correlate with the high gene expression level, we further performed whole genome sequencing in parallel with RNA-seq for the SARS-CoV-2 infected samples (fig. S9). The sequencing coverage was about 30X and more than 95% chimeric events were covered at least 10X, however, no chimeric junction reads between human and SARS-CoV-2 could be found in bins in terms of the highest chimeric level (Fig. 3C) and the most frequency of chimeric genes (Fig. 3, D-F). Moreover, there was no reads aligned to SARS-CoV-2 reference genome in all genome sequencing data, indicating that the viral integration into host DNA fragments is most unlikely through reverse-transcription.

Since the chimeric events truly existed in RNA-seq and showed preference to highly expressed genes, we speculate that they were likely caused by random priming between RNA templates in synthesizing first-strand and second-strand cDNA. However, we noticed that the chimeric loci showed random distributions, instead of preference to 5’ or 3’ termini, along both viral genome and host genes (Fig. 1D, and fig. S2C and S4A). It seemed that the original viral and host RNAs for chimeric genes were first cleaved into fragments which then formed the chimeric events between SARS-CoV-2 and host. In such a case, it is important to clarify sources of the chimeric events from either the digested RNA fragments or fragmentation during library construction.

To verify this speculation, we constructed the sequencing library with mixed RNA samples from SARS-CoV-2 infected Huh-7 cells and normal zebrafish embryos (Huh-7 RNA : Zebrafish RNA = 7 : 3), and then performed RNA-seq for this mixed library. With the same analysis pipelines, we found that the observed proportion of reads aligned to different species were similar as expected (fig. S10A). Intriguingly, the chimeric events were not only identified between SARS-CoV-2 and Huh-7 but also between SARS-CoV-2 and zebrafish, although most chimeric events were contributed by only one read (fig. S10B). Moreover, we consistently observed that both of viral chimeric genes in Huh-7 and zebrafish showed preference to the highly expressed genes (fig. S10, C-E), and the ratios of viral chimeric reads in human versus zebrafish was proportionally correlated to the ratio of read number of human to zebrafish (Fig. 4A). For viral loci, chimeric reads for both human and zebrafish also displayed same distribution and were enriched in N sub-genomic RNA (Fig. 4B). All these results indicate that the chimeric events of SARS-CoV-2 and host cells from RNA-seq were false-positive and mainly emerged during library constructions. We then performed regression analysis for chimeric level with gene expression levels using CPM or RPKM (normalizing gene expression with gene length) (Fig. 4C and fig. S11A). Although both methods produced positive correlations, the regression parameters by gene CPM were almost equal for human and zebrafish (Fig. 4C), suggesting the chimeric level were mainly related to the fragments in library. Additionally, we also found some reads representing the chimeric events between human and zebrafish (Fig. S11B), further supporting that the chimeric events artificially occur during library construction.

**Fig. 4.**
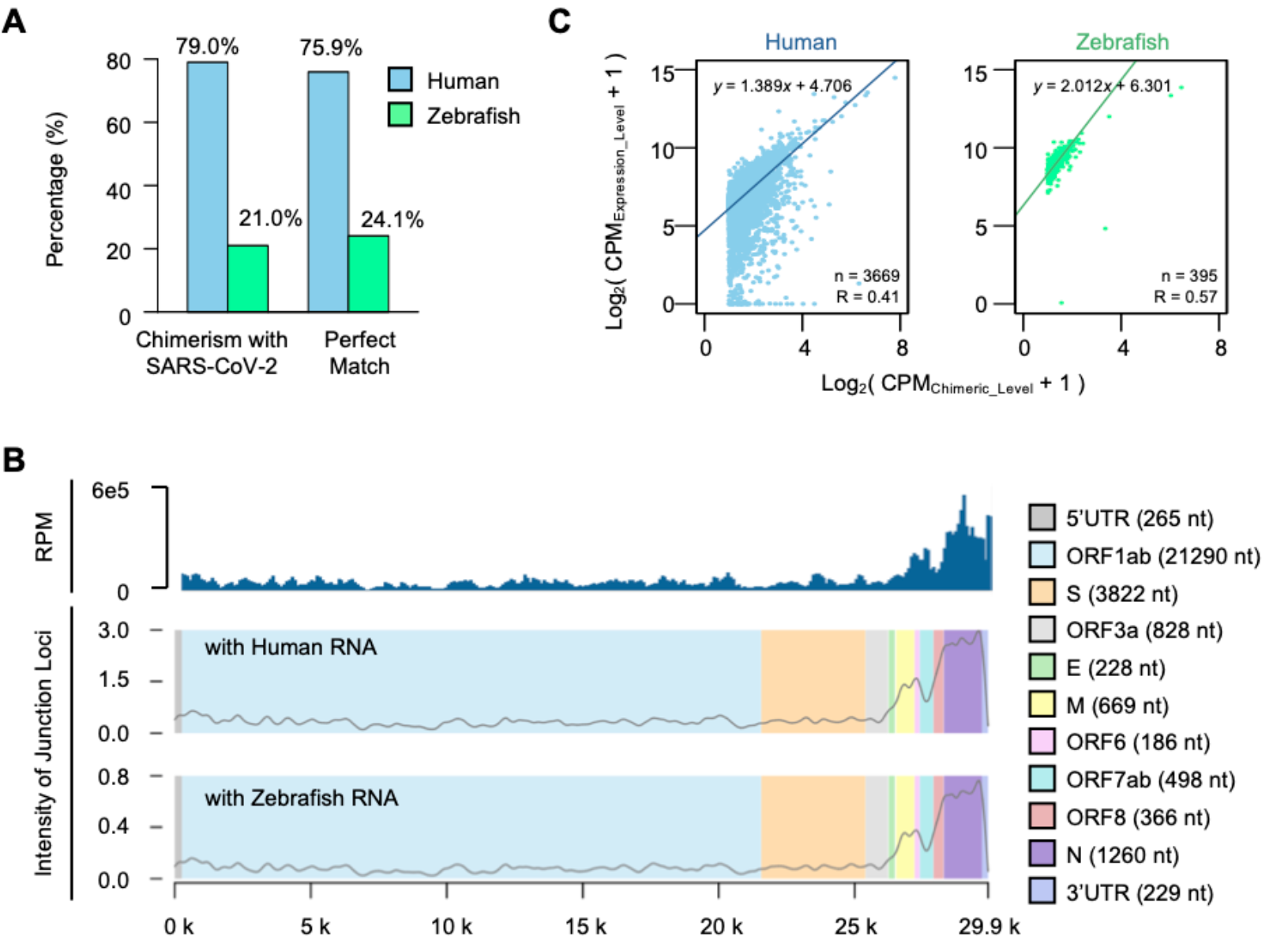
Chimeric reads in RNA-seq are falsely generated during library construction. (**A**) Proportions of viral chimeric reads with human (blue) and zebrafish RNA (green) were shown in left, while the corresponding ratio of sequencing depth for human and zebrafish, which defined as number of perfect matched reads, were shown in right. (**B**) IGV tracks displaying the depth of viral RNA along SARS-CoV-2 genome (top) and intensity of chimeric reads with human and zebrafish, individually (bottom). (**C**) Scatter plot displaying the correlation between chimeric level and counts of perfect matched fragments to chimeric genes for human (left) and zebrafish (right) from the same mixed library.

SARS-CoV-2 is highly contagious, and can cause severe clinical symptoms through infringing multi-organ systems (*21–26*), it is therefore necessary to clarify whether or not the viral RNA sequence could be integrated into the genome of patients leading to a long-term health risk. The recent report revealed the integrating events of SARS-CoV-2 with host genome in LINE-1 overexpressing human cells (*20*). To further clear this issue, we performed both RNA-seq and whole genome sequencing on SARS-CoV-2 infected human and monkey cells. Although we identified the presence of chimeric reads in RNA-seq, the analysis on bins with highest chimeric level and the most frequency of chimeric genes, in corroboration with the whole genome sequencing data, failed to show any chimeric reads (Fig. 3C-F). More importantly, chimeric reads were also identified in uninfected zebrafish embryos when mixing their RNAs with the RNAs from infected human cells at library construction step, therefore, providing the solid evidence that chimeric reads in viral infected samples are artificially introduced mainly through random ligations during library construction. In support, another team also observed the similar results based on their RNA-seq analysis (*27*). Although Zhang et al. found that SARS-CoV-2 could be reverse-transcribed and integrated into the genome of cultured cells overexpressing LINE-1 (*20*), it might be mainly due to the activation of retrotransposon but not natural characters of SARS-CoV-2. Overall our findings provide clear evidence that SARS-CoV-2 does not integrate into host genome, which will certainly help to alleviate the public concern about this issue.

## Author contributions

Y.-G.Y. and X.P. conceived this project, supervised the study; S.L. prepared the cells and virus infection with the help from T.D., Y.-N. Z., Y.Z., Y.Z. and P.L.; Y.Y., W.-J.L. and Y.-Q.S. performed the library construction experiments; Y.-S.C. and B.Z. performed bioinformatics analysis with the help from M.L.; Y.-G.Y., X.P., Y.Y., Y.-S.C., S.L., B.Z. and Y.-L.Z. discussed and integrated the data, wrote the manuscript. All authors read and approved the final manuscript.

## Data and materials availability

The RNA-seq and whole genome sequencing data supporting the conclusions of this article has been deposited in the Genome Sequence Archive under accession number subCRA005621 linked to the project PRJCA004871.

## SUPPLEMENTARY MATERIALS

Materials and Methods

Supplementary Text

Figs. S1 to S11

Tables S1 to S6

## Materials and Methods

### Cell culture and virus infection

HEK293T (Cell Resource Center, Institute of Basic Medicine Chinese Academy of Medical Sciences, 3111C0001CCC000091), Huh-7 (Cell Resource Center, Institute of Basic Medicine Chinese Academy of Medical Sciences, 3111C0001CCC000679), Calu-3 (Cell Resource Center, Institute of Basic Medicine Chinese Academy of Medical Sciences, 3111C0001CCC000032), Cos-7 (Kunming Cell Bank, Chinese Academy of Sciences, 3153C0001000000037) and MA-104 (Chinese Typical Culture Preservation Center Cell Bank, 3142C0001000000041) cells were cultured in DMEM (ThermoFisher,0030034DJ) supplemented with 10% fetal bovine serum (Gibco, 10099-141C) and penicillin-streptomycin (ThermoFisher,15140122) at 37°C, 5% CO_2_.

Cells were infected with SARS-CoV-2 (CDC of Guangdong province, GD108#), at a multiplicity of infection (MOI) of 0.1, and were collected for RNA isolation 24 hours post virus infection. All experiments with the SARS-CoV-2 virus were performed in the BSL-4 laboratory.

### RNA purification, library construction and sequencing

Total RNA from SARS-CoV-2 infected cell samples was extracted with Trizol reagent (Invitrogen, 15596026), and then subjected to the rRNA depletion with the Ribo-off rRNA Depletion Kit (Human/Mouse/Rat) (Vazyme, N406-02) following the manufacturer’s instructions. Libraries were constructed using the KAPA RNA HyperPrep Kit (KAPA Biosystems, KK8541) following the manufacturer’s instructions. Sequencing was performed on Illumina NovaSeq 6000 system with paired end 150 bp read length. Meanwhile, mixed sample of SARS-CoV-2 infected Huh-7 cells with zebrafish embryonic RNA was also used to prepare RNA-seq libraries, serving as the artificial chimeric RNA-seq reads control. For each infected sample, two replicates were performed, while three replicates were performed for the mixed sample.

### Genomic DNA isolation, library construction and whole genome sequencing

DNA samples were isolated by Universal Genomic DNA Kit (CWBIO, CW2298) according to the manufacturer’s instructions. The optical density values at 260/280 were approximately 1.6∼1.8. Genomic DNA of the same SARS-CoV-2 infected cell lines was prepared and whole genome shotgun libraries were constructed by using TruePrep DNA library Prep kit V2 for Illumina (Vazyme, Cat No. TD501-TD503), followed by sequencing on Illumina NovaSeq 6000 with paired end 150 bp read length. For each infected sample, two replicates were performed. About 100 Gb data were obtained for each replicate (**Supplementary Table S1**).

### Bioinformatics analysis of RNA-seq data

Raw FASTQ reads was processed to filter low quality bases and cut adapter sequence by fastp (version 0.20.1)^1^. The clean RNA-seq reads from SARS-CoV-2 infected cell were aligned to host genome appending with SARS-CoV-2 genome (NC_045512) by using STAR (version 2.7.7a, parameters as followed: --chimOutType Junctions SeparateSAMold WithinBAM HardClip -- chimSegmentMin 50 --chimScoreJunctionNonGTAG 0 --alignSJstitchMismatchNmax -1 -1 -1 -1 --chimJunctionOverhangMin 50 --outSAMtype BAM SortedByCoordinate --quantMode TranscriptomeSAM GeneCounts)^2^. The versions of genomes include GRCh38 for human (annotation release 102), Vero_WHO_p1.0 for green monkey (NCBI Chlorocebus sabaeus annotation release 102) and Mmul_10 for rhesus monkey (NCBI Macaca mulatta annotation release 103), and GRCz11 for zebrafish (NCBI: GCA_000002035.4). The duplicates were discarded. For RNA-seq reads from the mixed libraries of SARS-CoV-2 infected cells and uninfected zebrafish embryos were also aligned to human genome, zebrafish genome and SARS-CoV-2 genome with the same pipeline. The chimeric reads were also gained between each two genomes.

### Bioinformatics analysis of whole genome sequencing data

Reads from whole genome sequencing data were mapped to human genome plus SARS-CoV-2 genome by using BWA (version 0.7.17)^3^, and genome coverage statistics was obtained by samtools (version 1.12)^4^.

### Gene expression analysis

Gene expression level were determined by RPKM (Reads Per Kilobase per Million mapped reads) and reads counts of genes were calculated by cufflinks^5^. The expressed genes were then filtered by RPKM ≥ 1. To note, in **Fig. 4C**, the expression levels were defined as CPM without normalization of transcript length to represent the fragment depth of each transcript, directly. The *p* value for difference of expressions between chimeric genes and non-chimeric genes were calculated by Mann-Whitney U test (**Supplementary Table S2, S4 and S6**).

### Chimeric gene analysis

Chimeric reads between host and SARS-CoV-2 were obtained *via* the above STAR analysis pipeline. For these reads, the fragments from host genome were annotated by corresponding genome annotation reference by bedtools intersectBed^6^. The chimeric level was then defined as CPM (Counts Per Million reads), which were calculated by all chimeric reads from same gene and normalized by viral reads depth but not sequencing depth, except for results in Supplementary **Fig. S1**. The chimeric genes were then filtered by CPM ≥ 1 (**Supplementary Table S3, S5 and S6**).

The chimeric events were defined by loci of junctions between host and SARS-CoV-2. Chimeric events at different resolutions for (10 nt, 50 nt and 100 nt) were re-defined by corresponding steps, which were obtained by sliding windows at various resolution along host and viral genome.

### Analysis for accumulated expression and chimeric levels

The accumulated expression and chimeric levels were defined by sum of genes in related bins. Genes were first sorted in order of loci along each chromosome and then separated into one bin for each 100 genes. Those genes in the terminal of chromosome were defined as one bin though the number of genes might be less than 100. The accumulated expression level and chimeric level of each bin were calculated by sums of expression level and chimeric level from genes in the bin, respectively.

### Statistics and reproducibility

Non-parametric Mann-Whitney U-test (Wilcoxon rank-sum test, two-sided) is applied for calculating *p* value to assess the statistical significance of differences between two groups, which has also been mentioned in the related figure legends. Linear regression model is performed to evaluate the correlation between chimeric level and expression level (both RPKM and CPM).

**Fig. S1.**
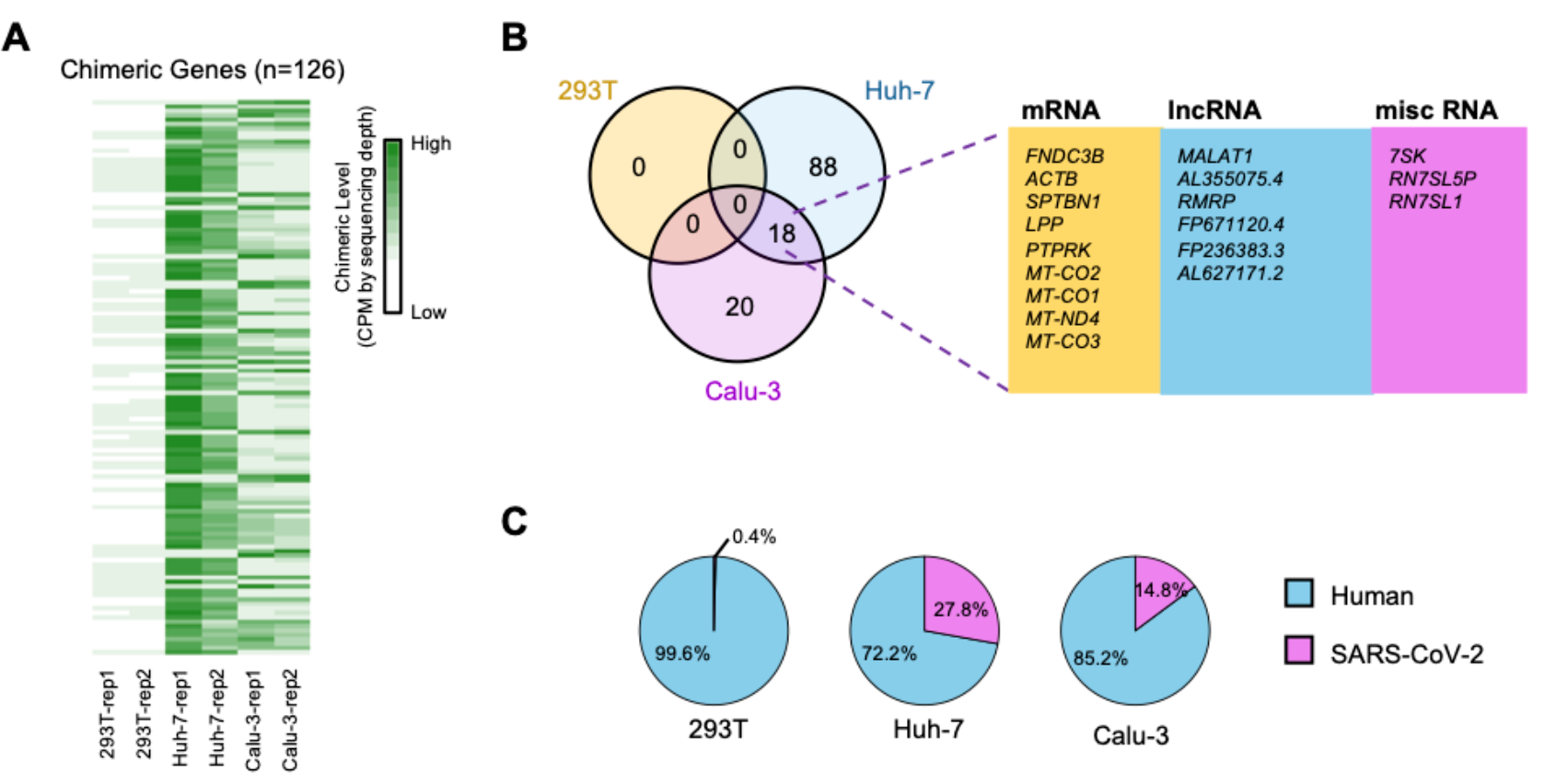
Analysis of chimeric genes with normalization of sequencing depth. (**A**) Heatmap displaying the chimeric level for filtered chimeric genes in each sample. The chimeric level was defined as CPM of chimeric reads and normalized by sequencing depth. (**B**) Venn diagram showing the number of shared and specific chimeric genes from **(A)** among 3 human cell lines. The overlapped genes between Huh-7 and Calu-3 cells mainly encode mRNAs, lncRNAs and miscRNAs. (**C**) Pie charts showing the proportions of reads aligned to human (blue) and SARS-CoV-2 (pink) genomes in infected 293T, Huh-7 and Calu-3 cells.

**Fig. S2.**
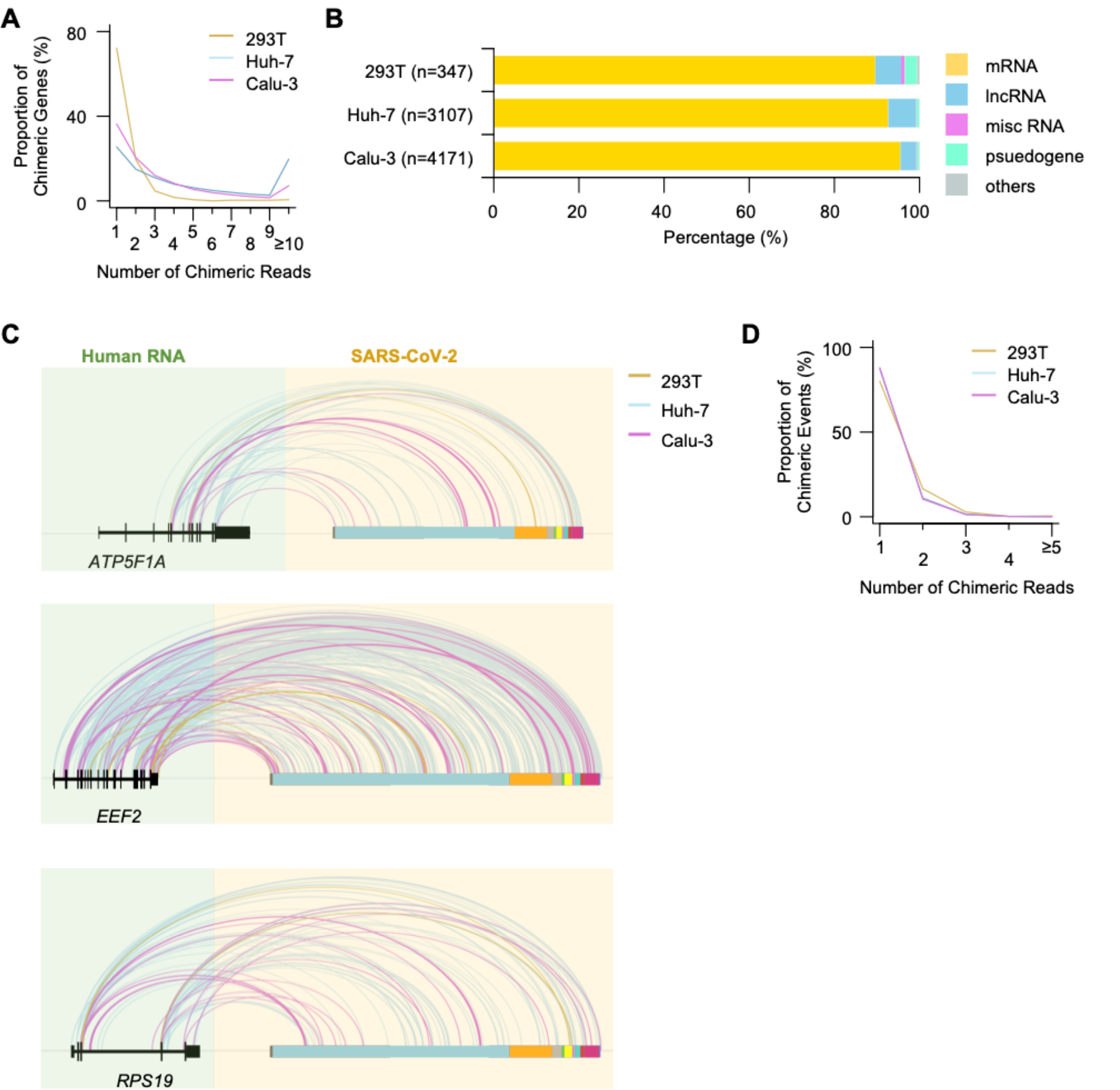
Viral chimeric gene transcripts were integrated with both mRNAs and non-coding RNAs from host. (**A**) Line chart displaying the proportion of chimeric genes supported by various numbers of chimeric reads in 293T (yellow), Huh-7 (blue) and Calu-3 (pink) cells. (**B**) Barplot displaying the proportions of RNA types for chimeric genes in 3 human cell lines. (**C**) IGV tracks displaying the junction loci in both human RNA and SARS-CoV-2 RNA for common chimeric gens *ATP5F1A* (top), *EEF2* (middle) and *RPS19* (bottom). The lines with different colors indicate the sources of chimeric reads. Yellow, blue and pink represents 293T, Huh-7 and Calu-3 cells, respectively. Blocks with different colors represent 5’UTR, ORF1ab, S, ORF3a, E, M, ORF6, ORF7ab, ORF8, N and 3’UTR along the SARS-CoV-2 genome from left to right. (**D**) Line chart displaying the proportions of chimeric events supported by various numbers of chimeric reads in 293T (yellow), Huh-7 (blue) and Calu-3 (pink) cells.

**Fig. S3.**
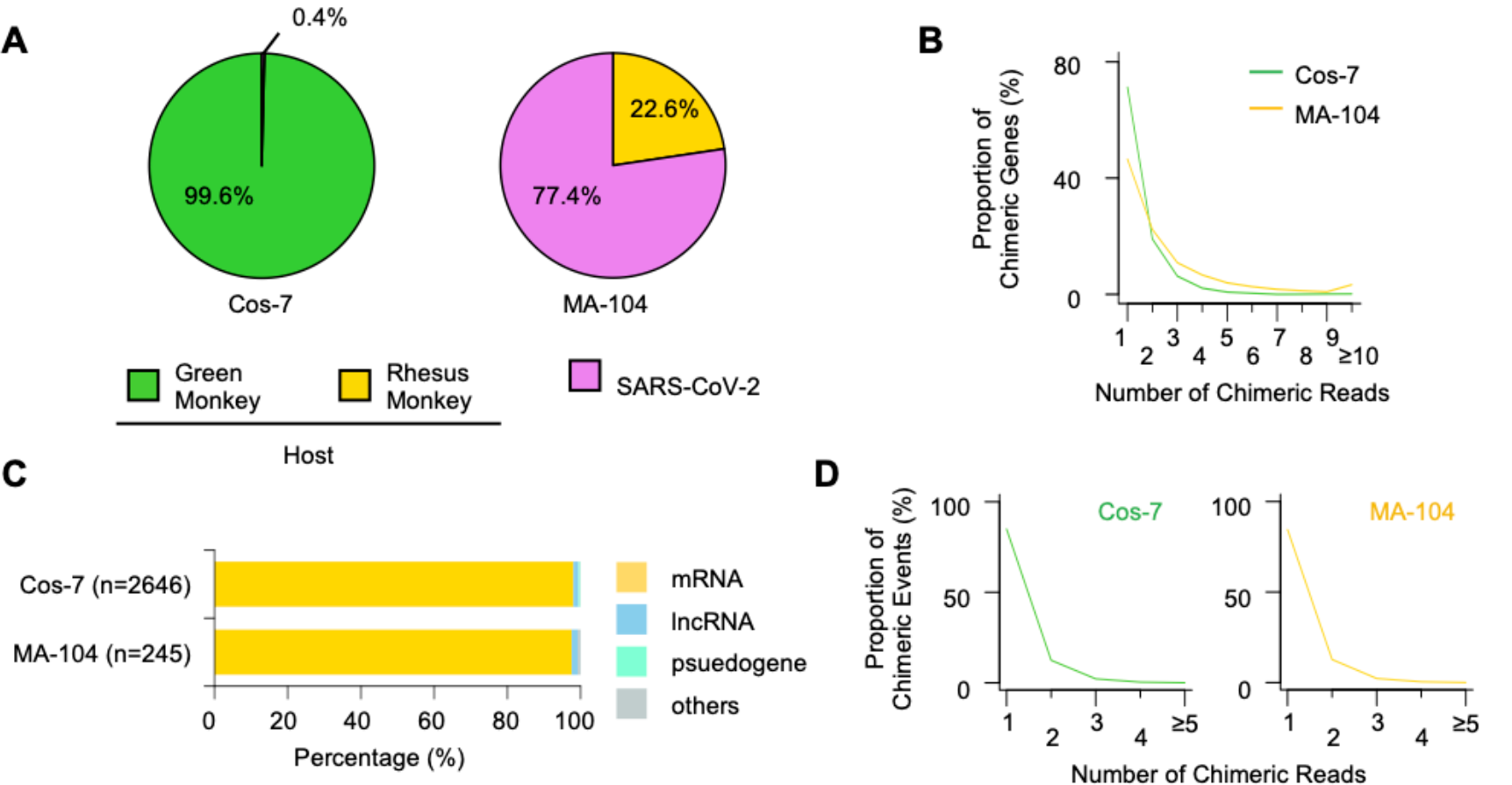
Analysis of chimeric genes in Cos-7 and MA-104 cells. (**A**) Pie charts showing the proportions of reads aligned to green monkey (green) and SARS-CoV-2 (pink) in infected Cos-7 cells (left), and proportions of reads aligned to rhesus monkey (yellow) and SARS-CoV-2 (pink) in infected MA-104 cells (right). (**B**) Line chart displaying the proportion of chimeric genes represented by various numbers of chimeric reads in Cos-7 (green) and MA-104 (yellow) cells. (**C**) Barplot displaying the proportion of RNA types for chimeric genes in Cos-7 and MA-104 cells. (**D**) Line chart displaying the proportions of chimeric events represented by various numbers of chimeric reads in Cos-7 (left) and MA-104 (right) cells.

**Fig. S4.**
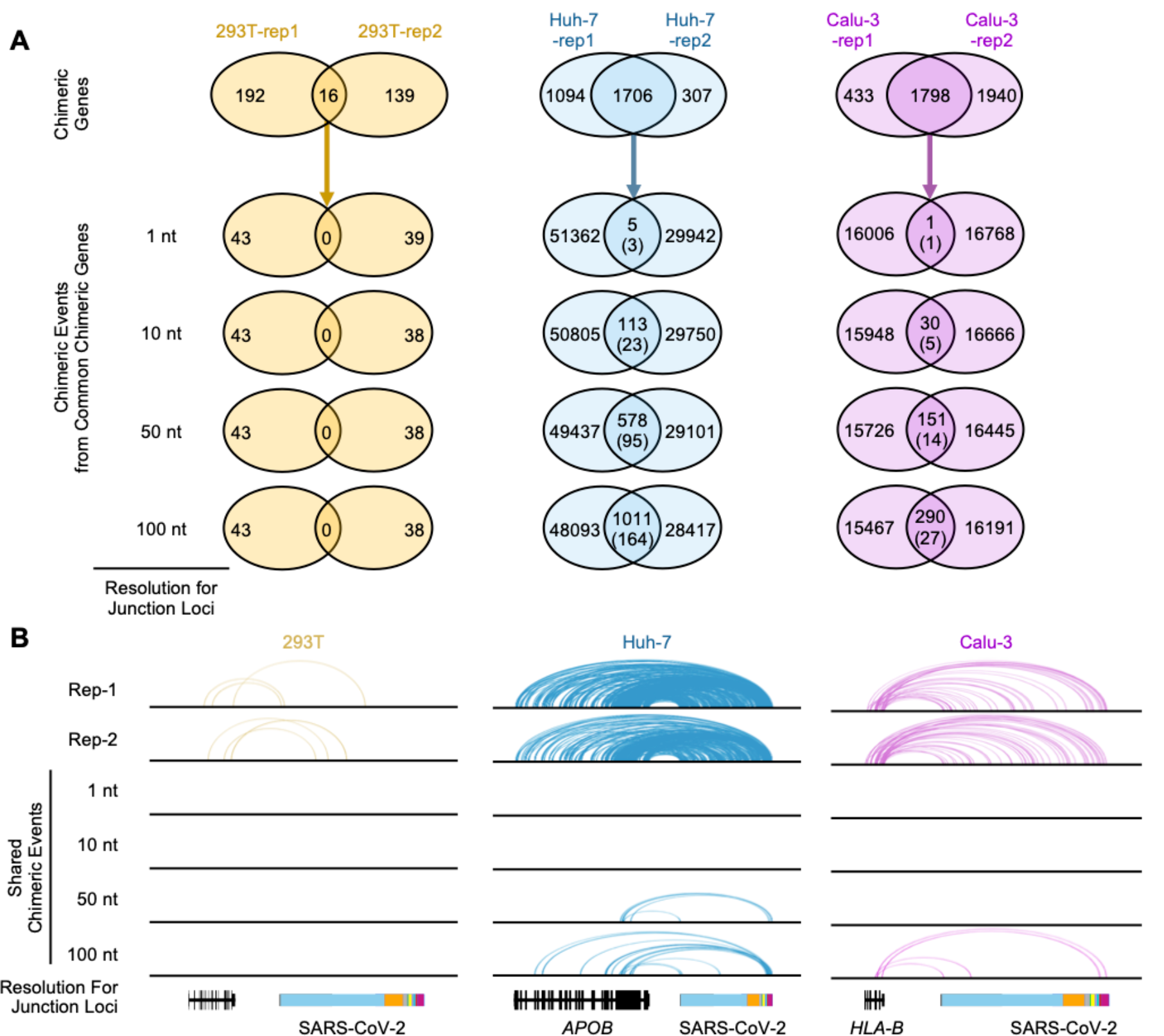
The chimeric events were not conserved between replicates. (**A**) Venn diagrams showing the number of shared chimeric genes for two replicates from same cell line (top) and conserved chimeric events on shared chimeric genes at different resolutions (1 nt, 10 nt, 50 nt and 100 nt). For venn diagrams of chimeric events, the values in brackets represents the numbers of chimeric genes containing the shared chimeric events. (**B**) IGV tracks displaying the junction loci in both human RNA and SARS-CoV-2 RNA for chimeric genes in each replicate (top). The shared chimeric events were displayed at different nucleotide resolutions (1 nt, 10 nt, 50 nt and 100 nt). Blocks with different colors represent 5’UTR, ORF1ab, S, ORF3a, E, M, ORF6, ORF7ab, ORF8, N and 3’UTR along the SARS-CoV-2 genome.

**Fig. S5.**
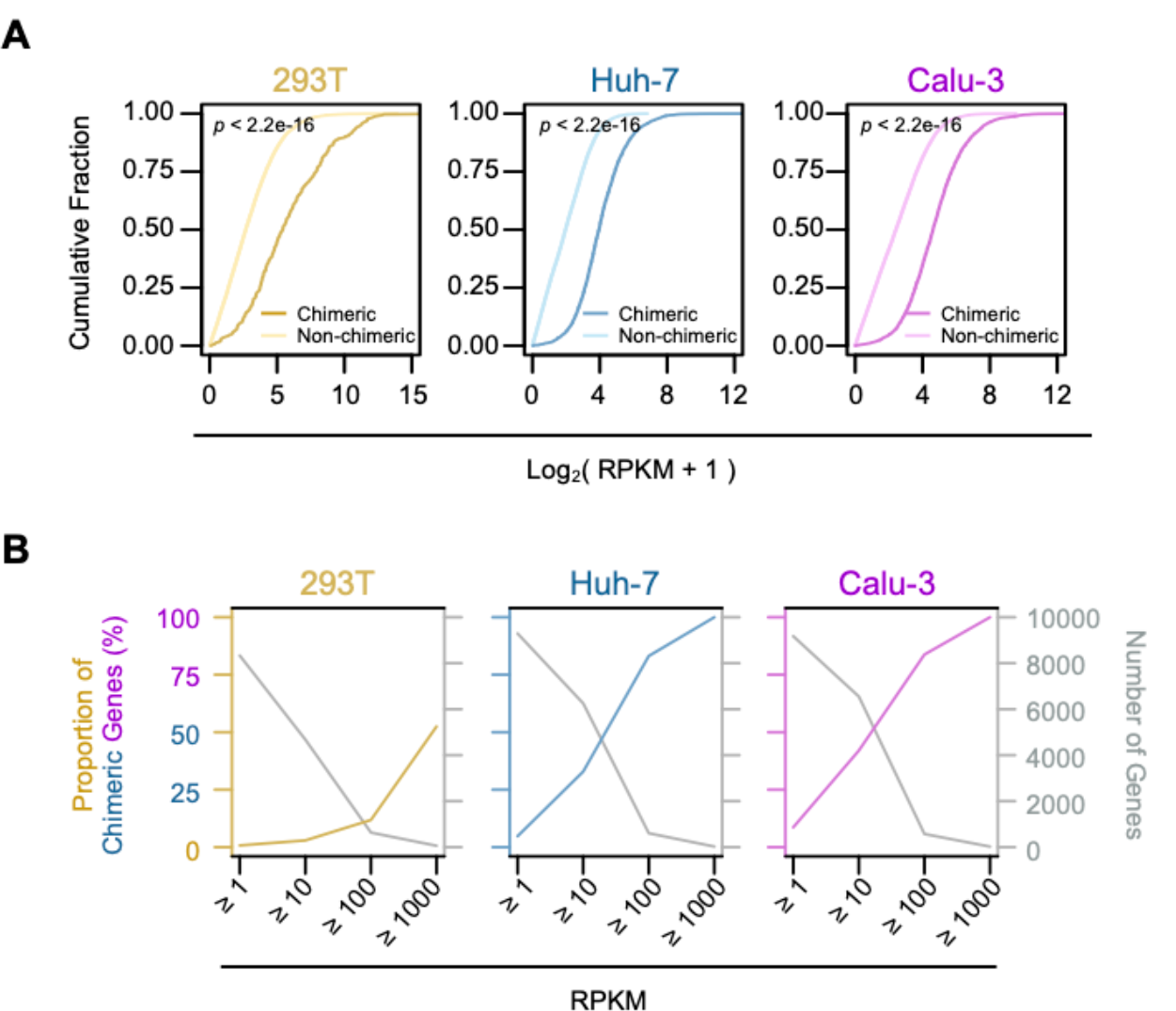
Chimeric genes were enriched in highly expressed genes. (**A**) Cumulative distributions of expression levels for chimeric genes (dark color) and non-chimeric gens (light color) in 293T, Huh-7 and Calu-3 cells. *P* values were calculated by Mann-Whitney U test. (**B**) Number of genes within different expression levels (grey) and proportion of chimeric genes in each pool within various expression levels. Yellow, blue and pink represent 293T, Huh-7 and Calu-3 cells, respectively.

**Fig. S6.**
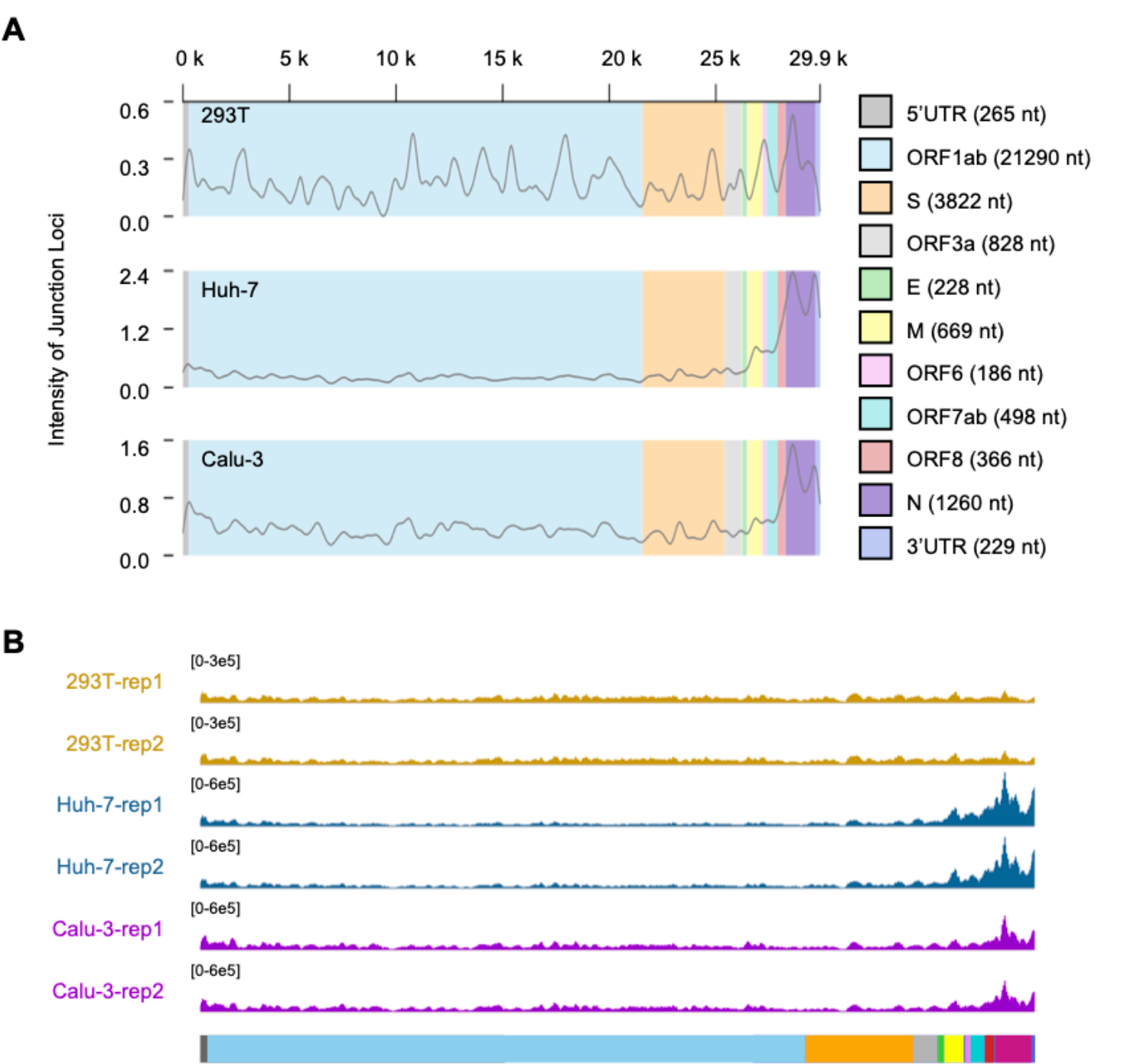
Chimeric intensity and sub-genomic RNA expression along SARS-CoV-2 genome. (**A**) IGV tracks displaying the intensity of chimeric reads along SARS-CoV-2 genome in infected 293T (top), Huh-7 (middle) and Calu-3 (bottom) cells. (**B**) IGV tracks displaying the depth of perfect matched reads along SARS-CoV-2 genome in two replicates of each cell line.

**Fig. S7.**
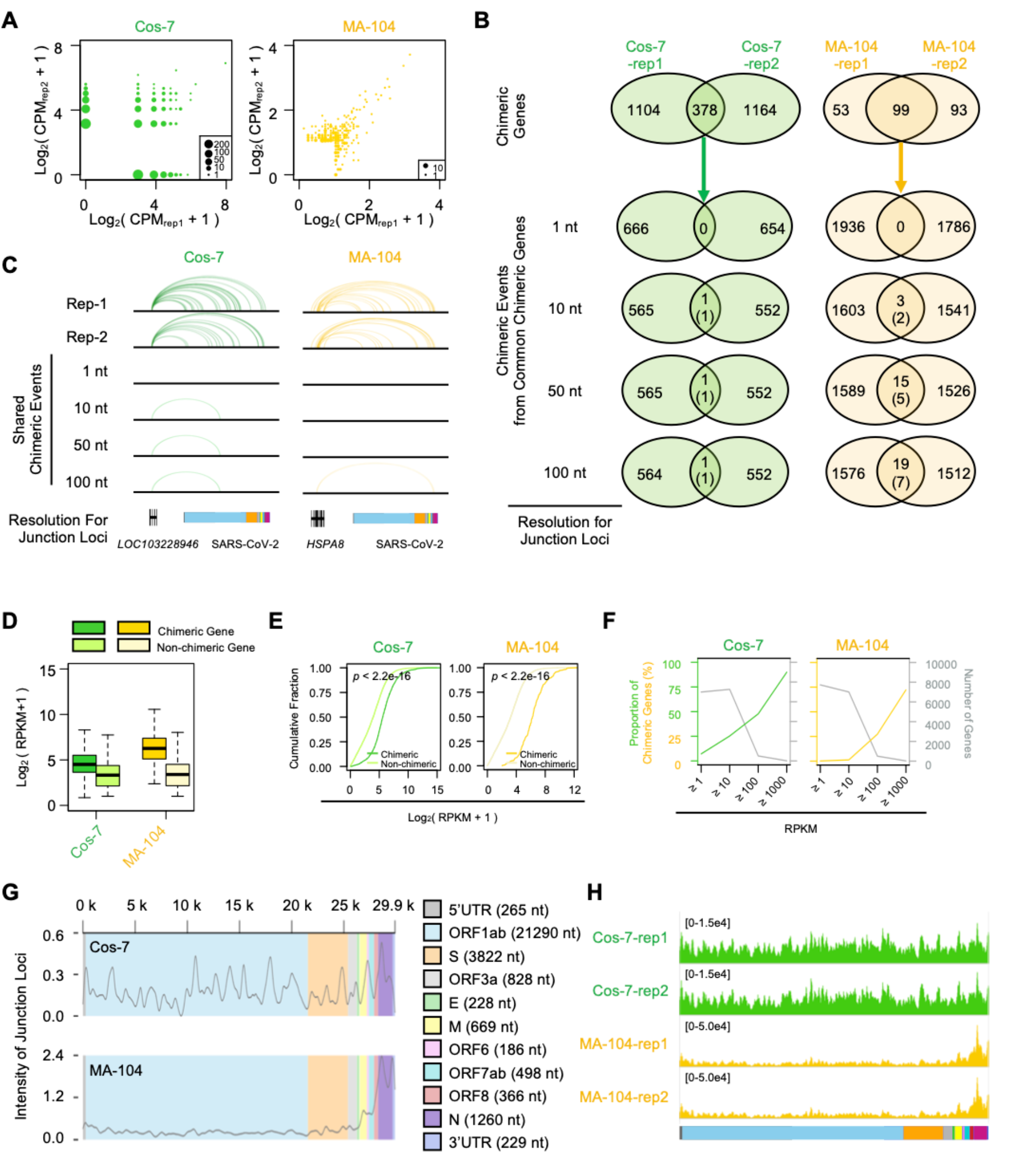
Both chimeric genes and events in Cos-7 (green monkey) and MA-104 (rehesus monkey) cells showed similar pattern to human. (**A**) Scatter plot displaying the chimeric levels for chimeric genes in two replicates. The sizes of scatters represent number of chimeric genes with the same distribution of chimeric levels in two replicates. (**B**) Venn diagrams showing the number of shared chimeric genes for two replicates of each cell line (top) and conserved chimeric events on shared chimeric genes at different nucleotide resolutions (1 nt, 10 nt, 50 nt and 100 nt). For venn diagrams of chimeric events, the values in brackets represent the numbers of chimeric genes containing the shared chimeric events. (**C**) IGV tracks displaying the junction loci in both monkey RNA and SARS-CoV-2 RNA for chimeric genes in two replicates (top). The shared chimeric events were displayed at different resolutions (1nt, 10nt, 50nt and 100nt). Blocks with different colors represent 5’UTR, ORF1ab, S, ORF3a, E, M, ORF6, ORF7ab, ORF8, N and 3’UTR along the SARS-CoV-2 genome. (**D**) Boxplot displaying the distributions of expression level for chimeric genes (dark color) and non-chimeric genes (light color) in infected Cos-7 and MA-104 cells. (**E**) Cumulative distributions of expression levels for chimeric genes (dark color) and non-chimeric gens (light color) in Cos-7 and MA-104 cells. *p* values were calculated by Mann-Whitney U test. (**F**) Number of genes within different expression levels (grey) and proportion of chimeric genes in each pool within varicose expression levels. (**G**) IGV tracks displaying the intensity of chimeric reads along SARS-CoV-2 genome in two replicates of infected Cos-7 (top) and MA-104 (bottom) cells. (**H**) IGV tracks displaying the depth of perfect matched reads along SARS-CoV-2 genome.

**Fig. S8.**
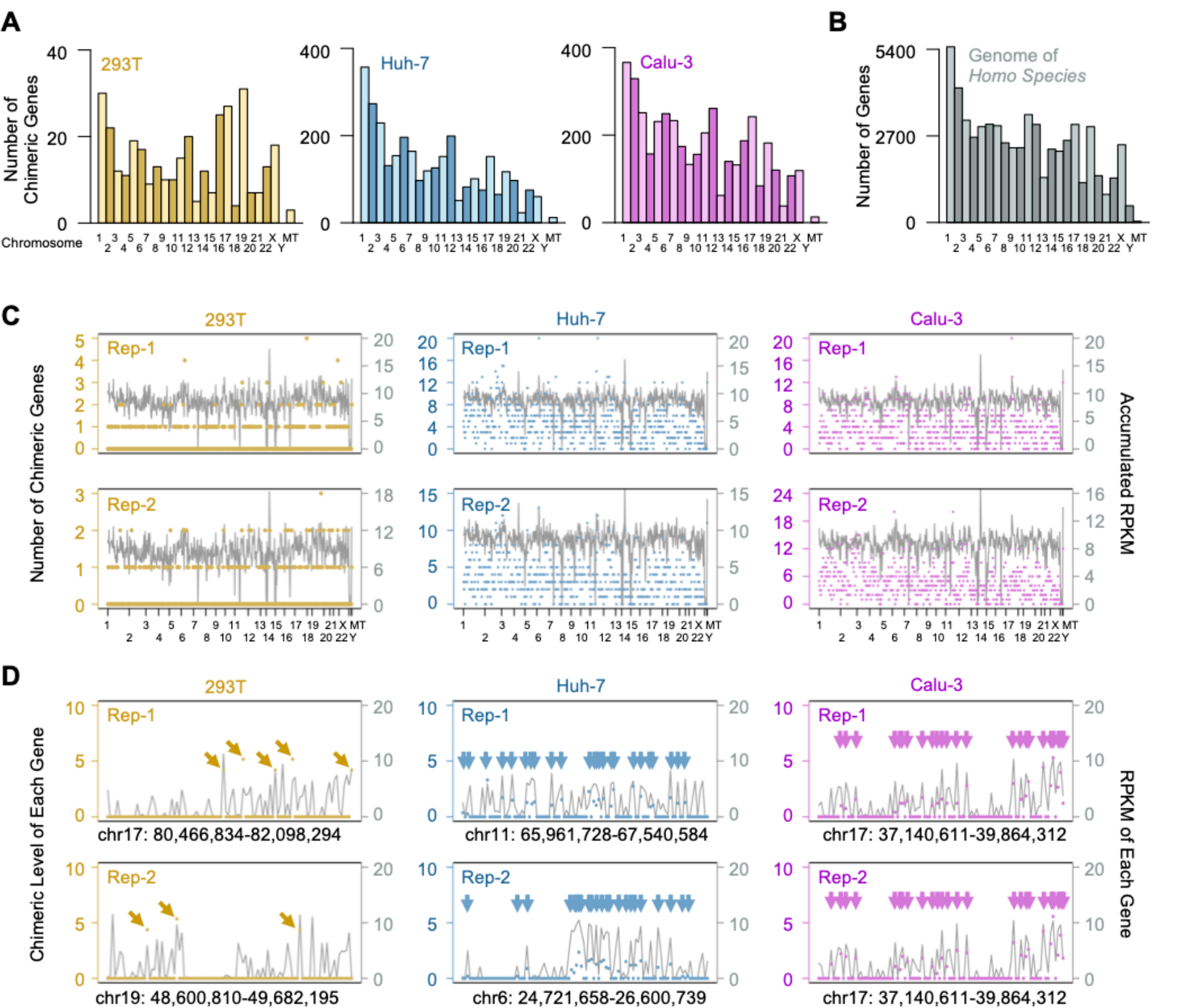
Information of chimeric genes along chromosomes. (**A**) Barplot displaying the numbers of chimeric genes along different chromosomes in infected 293T, Huh-7 and Calu-3 cells. (**B**) Barplot displaying the numbers of all annotated genes along different chromosomes. (**C**) The frequency and accumulated expression levels of chimeric genes in corresponding bins were displayed by scatters and lines, respectively. (**D**) Chimeric levels and corresponding expression levels of genes along the bins with the most chimeric genes were displayed by scatters and lines, respectively. The arrows indicate the loci of chimeric genes.

**Fig. S9.**
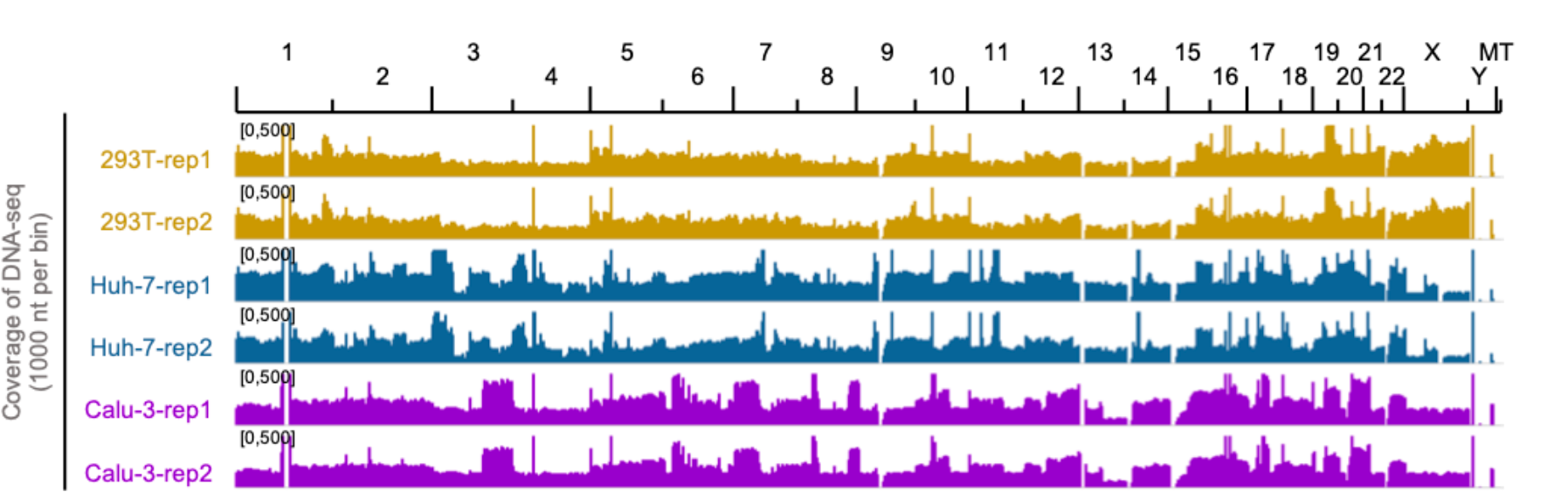
Coverage of reads from whole genome sequencing. IGV tracks showing the coverage of whole genome sequencing along genome in each sample.

**Fig. S10.**
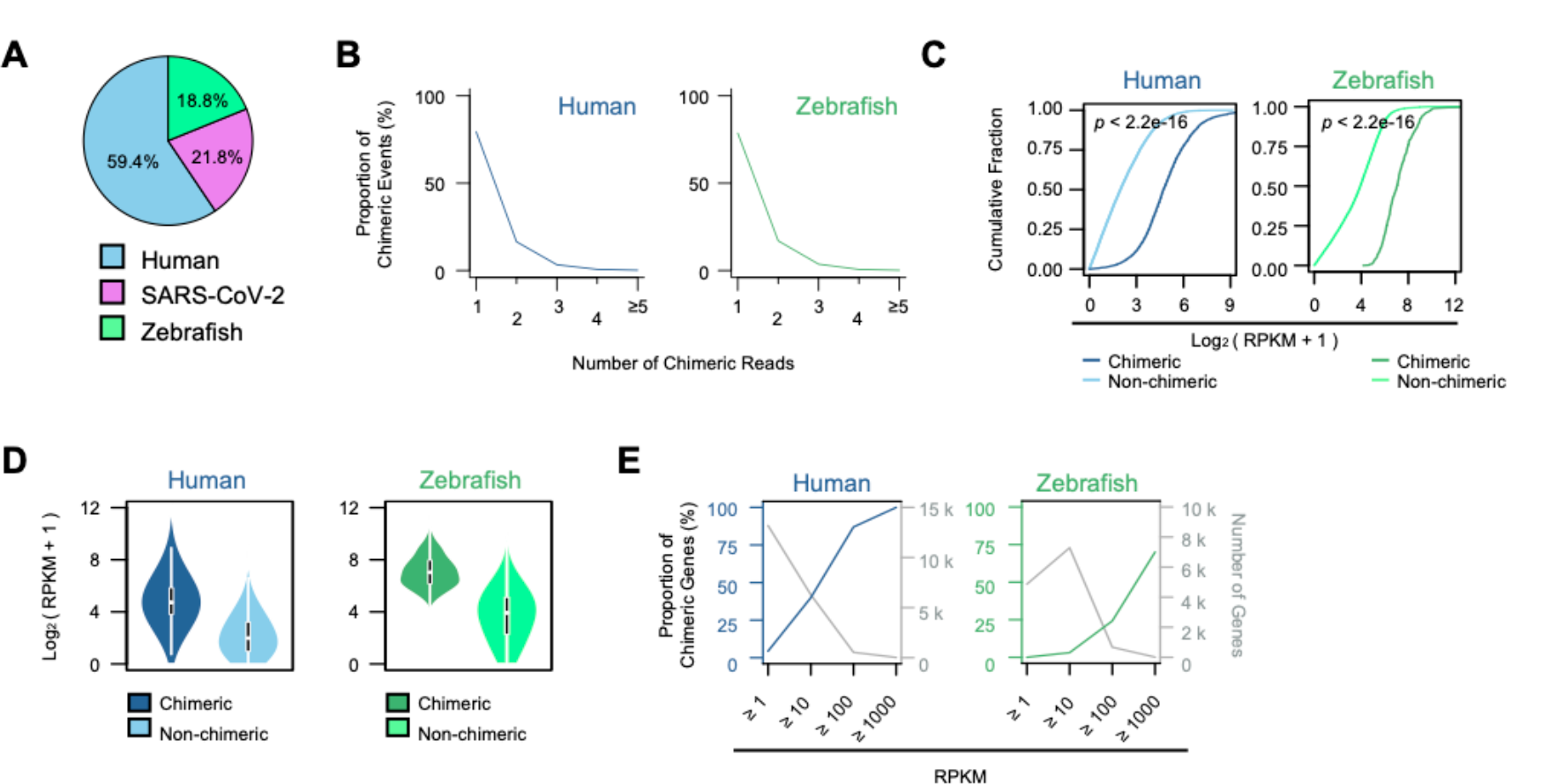
Chimeric genes could also be found in mixed RNAs from non-infected cells. (**A**) Pie chart showing the proportions of reads aligned to human (blue), SARS-CoV-2 (pink) and zebrafish (green) in the mixed library. (**B**) Line chart displaying the proportion of chimeric events represented by various numbers of chimeric reads in mixed library. Blue and green lines represent viral chimeric genes with human or zebrafish RNA, respectively. (**C**) Cumulative distributions of expression levels for chimeric genes (dark color) and non-chimeric gens (light color) for human (left) and zebrafish (right) RNAs. (**D**) Violin plot displaying the distributions of expression level for chimeric genes (dark color) and non-chimeric genes (light color) for human RNA (left) and zebrafish RNA (right). (**E**) Number of genes within different expression levels (grey) and proportion of chimeric genes in each pool within various expression levels. The left and right panels showing the human RNA and zebrafish RNA from the same mixed library, respectively.

**Fig. S11.**
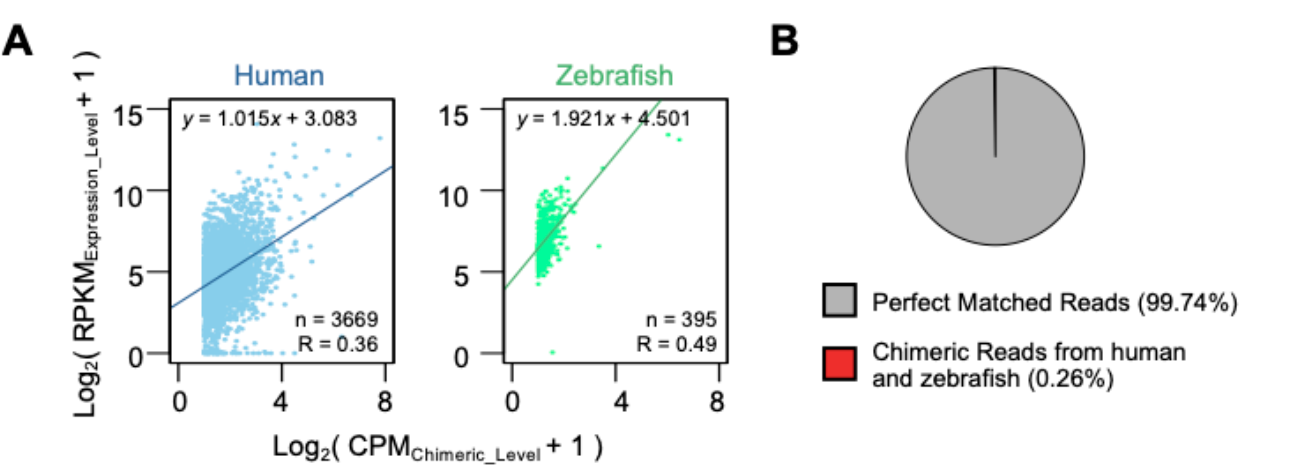
Chimeric events might come from library constructions but not natural events. (**A**) Scatter plot displaying the correlation between chimeric level and expression level of chimeric genes for human (left) and zebrafish (right) from the same mixed library. (**B**) Pie chart showing the proportions of reads identified as perfect matched reads (including reads aligned to human, zebrafish and SARS-CoV-2), and human-zebrafish chimeric reads (red).

**Table S1. Summary of RNA-seq and whole genome sequencing**.

**Table S2. Expression level of genes in infected human cell lines**.

**Table S3. Chimeric level of genes in infected human cell lines**.

**Table S4. Expression level of genes in infected monkey cell lines**.

**Table S5. Chimeric level of genes in infected monkey cell lines**.

**Table S6. Expression and chimeric level of genes from human and zebrafish in mixed RNA library**.

